# Sequencing Strategy to Ensure Accurate Plasmid Assembly

**DOI:** 10.1101/2024.03.25.586694

**Authors:** Sarah I. Hernandez, Casey-Tyler Berezin, Katie M. Miller, Samuel J. Peccoud, Jean Peccoud

## Abstract

Despite the wide use of plasmids in research and clinical production, the need to verify plasmid sequences is a bottleneck that is too often underestimated in the manufacturing process. Although sequencing platforms continue to improve, the method and assembly pipeline chosen still influence the final plasmid assembly sequence. Furthermore, few dedicated tools exist for plasmid assembly, especially for *de novo* assembly. Here, we evaluated short-read, long-read, and hybrid (both short and long reads) *de novo* assembly pipelines across three replicates of a 24-plasmid library. Consistent with previous characterizations of each sequencing technology, short-read assemblies had issues resolving GC-rich regions, and long-read assemblies commonly had small insertions and deletions, especially in repetitive regions. The hybrid approach facilitated the most accurate, consistent assembly generation and identified mutations relative to the reference sequence. Although Sanger sequencing can be used to verify specific regions, some GC-rich and repetitive regions were difficult to resolve using any method, suggesting that easily sequenced genetic parts should be prioritized in the design of new genetic constructs.

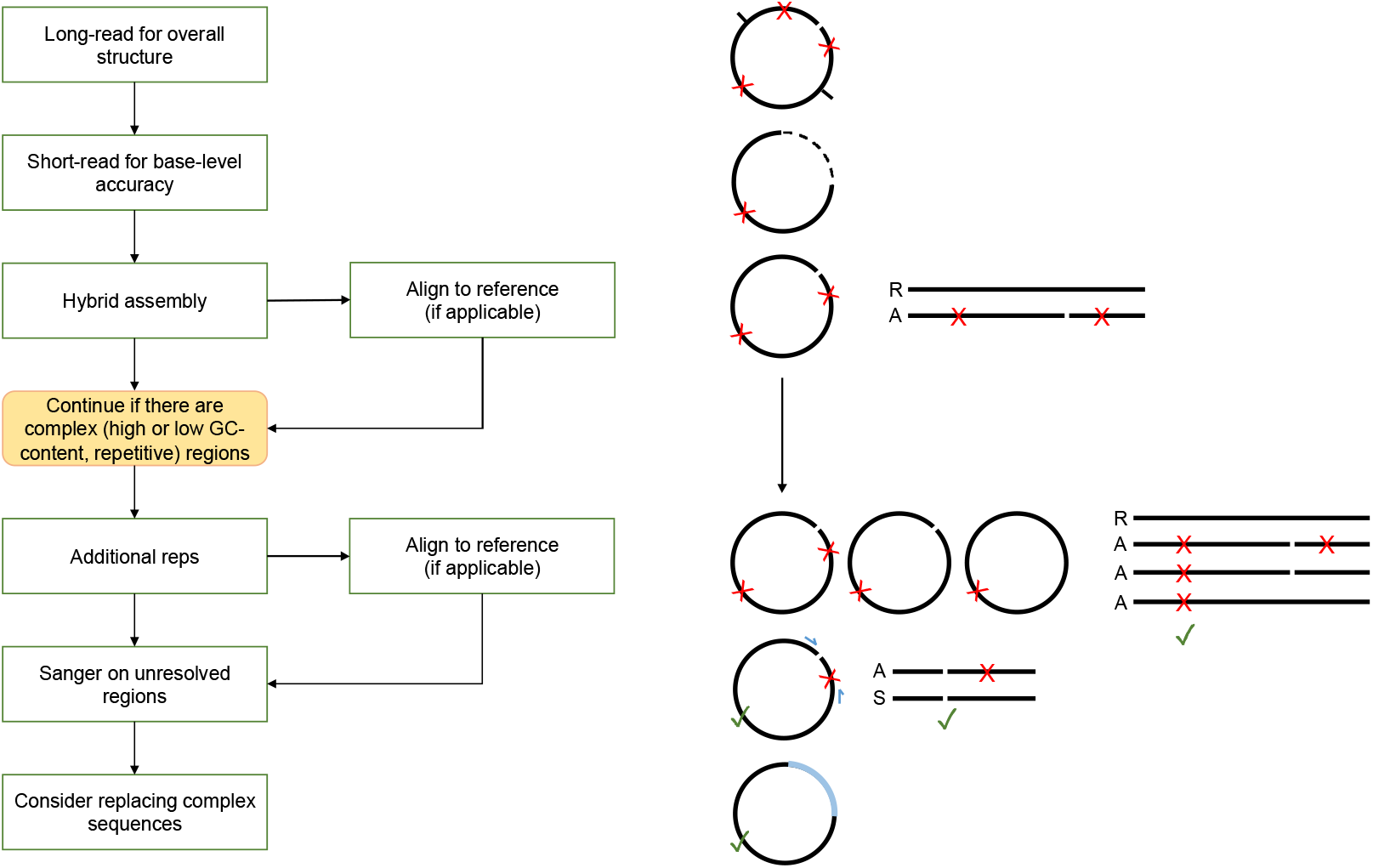
GRAPHICAL ABSTRACT.

## INTRODUCTION

Plasmids are critical tools used in research, industrial, and clinical settings for applications such as recombinant gene expression, designing genetic circuits, and the generation of clinical products like vaccines [1-8]. Sequence verification of these increasingly large plasmid libraries is critical to ensuring the expected product is made and to evaluate the biological effects of spontaneous and intentional mutations, including single nucleotide polymorphisms (SNPs) [2-5, 9]. Although DNA sequences are generally designed and documented digitally, and the necessity of openly providing DNA sequences and thorough gene annotations has been discussed many times, it is not uncommon for researchers to have only a vague plasmid map and/or no reference sequence [10-13]. Many plasmids are generated by inserting a gene of interest into a plasmid backbone, and sequence verification is often overlooked in favor of simpler “confirmation” methods, such as PCR amplification or restriction digests. Yet, without confirming a plasmid’s sequence, unrecognized deviations from an expected sequence threaten the accuracy of biological insights gained using such a plasmid.

Compounding a reluctance to perform sequence verification is the lack of dedicated plasmid assembly tools. Amidst the many tools for genome and metagenome assembly, there are some designed to identify plasmids in the assembly of these larger datasets [14-22]. However, many of these methods can struggle to accurately reconstruct small (<25 kbp) plasmids or miss them altogether [23, 24]. To accelerate plasmid verification in high-throughput settings, assembly pipelines should avoid the need for a reference sequence or manual intervention, as required by some methods [3, 25]. *de novo* sequence assembly is preferred to reference-based assembly as it can also help overcome reference bias and identify unexpected mutations [1, 2, 11, 26-28]. Here we used the two major end-to-end *de novo* assembly pipelines designed for plasmids: Epi2ME, a tool from Oxford Nanopore Technologies (ONT) for long-read assembly, and the newly developed open-source plasmid tool, PlasCAT, which can perform short-read, long-read, and hybrid assembly [2, 29-31].

Although Sanger sequencing has long been the gold standard for sequencing, it requires a reference sequence to design primers and is limited to short (∼800 bp) sequences, necessitating many reactions to verify a whole plasmid [32-36]. The need to sequence unknown DNA templates led to the introduction of next-generation sequencing (NGS) technology, namely the Illumina fragmentation-based approach [5, 32-34, 37]. While the short-read sequencing fragments (∼250 bp) generally provide good template coverage and high sequence accuracy, biases introduced in PCR steps result in the underrepresentation of GC-rich, GC-poor and repetitive regions [32, 33, 38-41]. Short-read sequencing can also underperform with low diversity libraries [38, 40]. Thus, third-generation sequencing methods that allow the sequencing of reads that are thousands of nucleotides long – as long as a plasmid itself – have been developed [3, 34]. These long-read sequencing methods can be faster and cheaper than fragmentation-based approaches, and better resolve long, complex sequences, but has historically had lower accuracy than its predecessors and can struggle with smaller templates [1, 2, 24, 42-44]. Recent advancements in genome assembly tools have indicated that a hybrid approach, using both short and long reads, can produce improved assemblies [20, 24, 43, 45]. Nevertheless, the ability to sequence growing libraries of DNA sequences, including genomes, with base-level precision is a continued pursuit [46]. To our knowledge, a hybrid approach to *de novo* plasmid assembly has not yet been systematically interrogated.

Here, we evaluated the ability of the short– and long-read sequencing methods available from Illumina and ONT, respectively, to generate accurate *de novo* assemblies of plasmid sequences. We found that a hybrid assembly approach, using both short and long reads, produced the best assemblies. The short-read assemblies were limited by the quality and quantity of DNA used and struggled to assemble GC-rich regions, whereas the long-read assemblies had a higher incidence of insertions and deletions (collectively, indels) and mutations, as has been previously suggested [23, 43, 47]. We used Sanger sequencing to confirm several discrepancies between the assembly sequences and the plasmid reference sequences and found several cases where the assemblies consistently differed from the expected reference sequence. Importantly, *de novo* assembly outperformed reference-based assembly, which frequently showed reference bias and often did not match Sanger data. Thus, *de novo* hybrid assembly is the preferred method for high-throughput plasmid sequencing.

## MATERIALS AND METHODS

### Reagents

The Zyppy-96 Plasmid MagBead Miniprep Kit was purchased from Zymo Research (Irving, CA, USA, #D4102). The long-read library preparation kit and R10.4 flow cell were purchased from Oxford Nanopore (UK, #SQK-RBK114.96 and #FLO-MIN114). For the short-reads, the ILMN DNA LP (M) Tagmentation 96 library preparation kit, MiSeq cartridges, and iSeq cartridges were purchased from Illumina (San Diego, CA, USA. #20060059, #MS-103-1003, and #20031374 respectively). Sanger sequencing primers were designed and ordered from IDT (Coralville, IA). The BigDye Terminator™ V3.1 and BigDyeXTerminator™ purification kits were both purchased from ThermoFisher (Waltham, MA. USA, #4337454 and #4376486 respectively).

### Biological Resources

The plasmids used for library preparation, sequencing, and analysis were obtained from three vendors. Twelve plasmids were synthesized and sequence-verified by Twist Biosciences (San Francisco, CA), and eleven were procured from Addgene (Watertown, MA, USA). One plasmid solution was taken from a transfection kit where the mixture contains two plasmids with the same vector backbone, roughly 6000 bp, and two different inserts, around 1350 bp and 650 bp (Gibco #A14635) [48, 49]. The array of plasmids was given unique identifiers (e.g., Plasmid 1234) to anonymize the data set. This allowed for true *de novo* assembly, where the different methods could be compared on overall accuracy among the generated datasets.

### Plasmid Isolation and Sequencing

#### Plasmid Isolation

The plasmid DNA was extracted from each of the 24 isolates on an epMotion 5075 TC liquid handler (Eppendorf, Hamburg, DE) using the Zyppy-96 Plasmid MagBead Miniprep Kit (Zymo Research, Irving CA, USA), according to manufacturer’s instructions with the modification of pipet mixing during the lysis and neutralization steps and an extended elution time of 10 minutes. Samples were all quantified on a Synergy LX plate reader to determine the quality and quantity of samples after extraction via miniprep. All samples were required to have at least 35 ng/uL and an A260/280 purity reading of >1.8.

#### Oxford Nanopore Sequencing

Post isolation, 50 ng of each isolate was used for sequencing with the MinION Sequencer (Oxford Nanopore, UK). These sequencing libraries were prepared using the Rapid Barcoding Kit (#SQK-RBK114.96) with the Flow Cell (#FLO-MIN114) according to the manufacturer’s instructions. Samples were run on the MinION with a maximum read length kept of 25 kbp. FASTQ files were generated from the super-high accuracy method of the Dorado basecaller within the MinKNOW software and were used for sequence validation, comparison, and evaluation.

#### Illumina Sequencing

After isolation, 200 ng of each isolate was used for sequencing on the MiSeq and iSeq (Illumina, CA). Both sequencing libraries were prepared on an epMotion 5075 TC liquid handler (Eppendorf, Hamburg, DE) using the ILMN DNA LP (M) Tagmentation 96 IPB kit protocol as described by the manufacturer. The pooled libraries were spiked with 1% v/v PhiX Control V3 (Illumina, San Diego, CA) and were diluted to a final loading concentration of 10 pM and 100 pM for the MiSeq and iSeq, respectively. The diluted libraries (600 µL and 20 µL) were loaded onto a MiSeq Reagent Nano Kit v2 (500-cycles) and iSeq 100 i1 Reagent v2 (300-cycle). FASTQ files generated were used for sequence validation, comparison, and evaluation.

#### Sanger Sequencing

Sequence validation was performed as needed for templates with generated sequence discrepancies. Primers were designed and ordered through IDT between 18 to 25 bps long to satisfy the following requirements: a GC content of 50% or higher, a melting temperature around 70 C, and no secondary structure (Supplementary Table S1). Fragments were prepared following the BigDye Terminator™ V3.1 kit as described, and the 10 µL reactions were diluted to 0.5X using the BigDyeXTerminator™ purification kit where described. Samples were sequenced using the LongFrag_BDX protocol. Generated results were immediately uploaded to SnapGene and compared to generated data.

#### *De Novo* Sequence Assembly

*De novo* sequence assembly was primarily performed using PlasCAT, an open-source plasmid assembly pipeline that was recently adapted to a web application (sequencing.genofab.com) [2, 29, 31]. In brief, the pipeline generates assemblies from short-reads, from long-reads, or from a hybrid approach (i.e. both short and long reads) via the gold-standard genome assembly tool, Unicycler [20]. The pipeline also performs some pre-processing of the data, either through Trimmomatic for short-reads [50] or Filtlong for long reads (https://github.com/rrwick/Filtlong), and subsets the long reads to a particular coverage using Rasusa [51, 52]. The short reads were filtered to a minimum length of 50 bp and a minimum quality score of 35. Filtlong was used to keep the best 80% of long reads (based on quality and length) and remove reads above the maximum read length of 20,000 bp [53]. Subsetting with rasusa was done using the default estimated size of 5,000 bp and 500X coverage, which gave better long-read assemblies than the default 1000X coverage. Long-read and hybrid assemblies are polished with racon [54]. Long-read assemblies were also generated using Oxford Nanopore’s EpiPI2ME platform to serve as a method comparison (https://labs.epi2me.io/). This pipeline uses Flye for long-read assembly [55], the Medaka polisher (https://github.com/nanoporetech/medaka) [30, 56], and Trycycler to generate a consensus assembly [25]. For Epi2ME, we used the default estimated size of 7000 bp, 60X coverage, and end trimming of 150 bp.

## RESULTS

### Plasmid Sequencing and Assembly Pipelines

We sequenced a set of 24 plasmids and performed *de novo* assembly using short-reads from the Illumina MiSeq and long-reads from the ONT MinION. Three technical replicates were sequenced from each plasmid, all pulling from the same initial purified plasmid solution. The open-source tool PlasCAT was used to generate assemblies in three ways: short-read only, long-read only, and a hybrid approach using short and long reads (Figure 1). This design allowed us to compare both the reproducibility of each sequencing technology across repeated library preparations and evaluate different plasmid assembly approaches.

**Figure 1.**
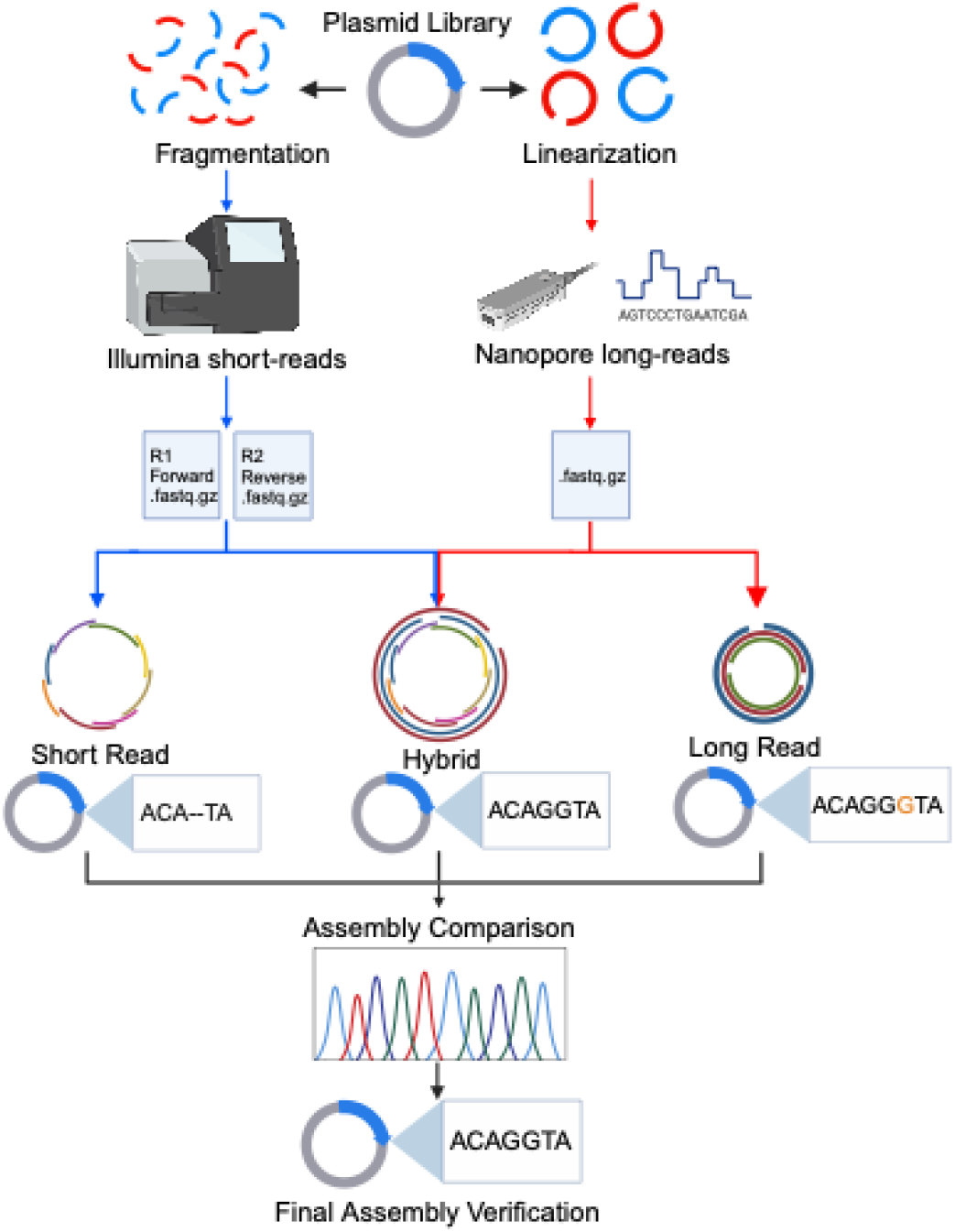
Overview of sequencing workflow. Multiple sequencing runs were performed on 24 plasmid samples. For short-read sequencing, plasmids were fragmented and chemically indexed before being loaded onto the Illumina MiSeq. Forward and reverse reads were generated and used with PlasCAT to generate both short-read and hybrid assemblies. For long-read sequencing, the plasmids were linearized, chemically indexed, and loaded onto the Oxford Nanopore MinION. FASTQ files were generated and used for both long-read and hybrid assemblies. Sanger sequencing was used to confirm regions with discrepancies between assemblies.

Of the 72 total samples prepared, all generated data on the Nanopore sequencer, while only 69 samples generated FASTQ files with data on the MiSeq. Of the three failures, two were replicates of the same sample. Issues with sample dropouts prohibited us from generating assemblies for these three samples and were attributed to the library preparation procedure and not the assembly process. The short reads were trimmed to maintain a per-base quality score of at least 35 and reads shorter than 50 bp were removed (Supplementary Figure S1). Before filtering, there were roughly 50,000 reads (combined forward and reverse) for each sample, representing at least 500 million bases per run. After filtering, only about 5,000 reads were retained per sample (Supplementary Figure S1). The long reads were filtered with Filtlong. There were initially about 50,000 reads per sample, representing over 3 billion bases per run. Filtlong retains the best 80% of the data, based on both quality and length. This is evidenced by an increase in average read length (from ∼4,000 to ∼6,000) and minimum read length (from 500 to ∼1,000 bp) in post-filtered samples, despite the decrease in maximum read length to 20,000 bp. After filtering, there were about 25,000 reads per sample (Supplementary Figure S1).

### Hybrid *De Novo* Plasmid Assembly Outperforms Short-Read or Long-Read Assembly

To quantify the robustness of each assembly method, we devised two scoring methods: an assembly score to represent the success of a particular assembly, and a sequence agreement score to assess the reproducibility of assemblies across multiple runs (Figure 2). An assembly score of 1 indicates a single contig was returned (a success), while a 0 was given if no assembly or if multiple contigs were returned (a failure). An overall assembly score was obtained by summing the scores of each of the three runs. A sequence agreement score of 1 indicates that the sequences of all successful (non-fragmented) assemblies were the same, or a 0 if not. Plasmids that only had one successful assembly were excluded from this scoring. The overall assembly and sequence agreement scores were converted into percentages based on the number of included runs and samples, respectively. We included one sample that was a mixture of two plasmids (Plasmid 3589) to see whether the plasmid assembly pipelines would generally return one contig, or if it could resolve mixtures of similar plasmids, but it was excluded from our formal data analysis.

**Figure 2.**
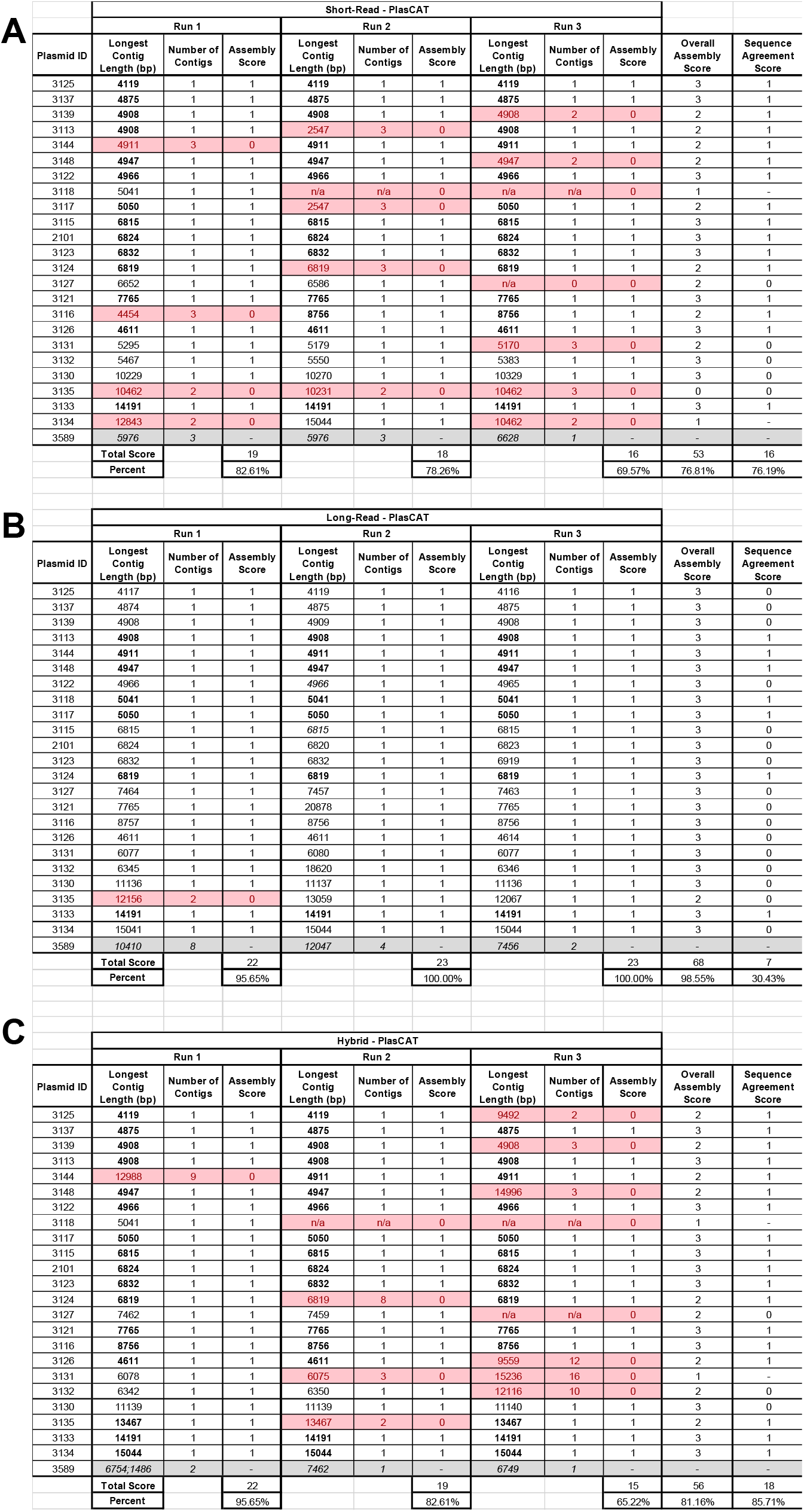
Hybrid assemblies outperform short– and long-read assemblies. *De novo* assemblies were generated from short-reads (A), long-reads (B), or from both (hybrid, C). All assembly pipelines produced some fragmented assemblies (>1 contig) which were considered failures (red). Only the length of the longest contig is reported in these cases. Some short-read sequencing preparations did not produce sufficient data for assembly (n/a, red). An assembly score of 1 indicates a successful assembly (non-fragmented), and these are summed across the three replicates to generate an overall assembly score (maximum of 3). A sequence agreement score of 1 indicates that all successful assemblies were exact matches for one another, and the corresponding assembly lengths are bolded. Samples with only one successful assembly were not given a sequence agreement score and were excluded from the percentage calculation. The sample containing a mixture of two plasmids (Plasmid 3589) was also excluded from this analysis, as it was not expected to return only one contig. The hybrid assemblies had the highest sequence agreement scores, followed by short-read and then long-read assemblies. The long-read assemblies had the highest overall assembly score but failed to produce high sequence agreement scores, indicating lower reproducibility of assembly results.

We generated short-read assemblies for all 69 of the samples that produced sufficient data. Most of the assemblies were single contigs, however the pipeline returned 13 assemblies with multiple contigs (Figure 2A). Given that we expected all samples to result in a single contig representing the plasmid, these 13 assemblies were considered failures and given an assembly score of 0. There was only one sample that received an overall assembly score of 0, for which all three assemblies were fragmented (Plasmid 3135). Nevertheless, nearly 77% of runs resulted in a successful assembly. Furthermore, 76% of samples had good sequence agreement scores, indicating that the assembly of most plasmids by short-read sequencing is reproducible across repeated library preparations. The sequence agreement score is ultimately more important than the assembly score, since the consistency provides researchers with a higher level of confidence that their assemblies are correct. Notably, there were 5 samples, including most of the plasmids larger than 10 kb, which received a sequence agreement score of 0 for their short-read assemblies and for which no consensus could be reached (Figure 2A).

We compared the assemblies generated from one sequencing run using the iSeq, which produces 151 bp reads, to the MiSeq, which produces 251 bp reads. Overall, the assemblies generated by the iSeq were similar to the assemblies from the MiSeq (Supplementary Table S2). Four of the iSeq assemblies contained multiple fragments, which was comparable to the failure rate of the MiSeq assemblies. Although the assemblies generated by the iSeq reads were not considerably better or worse, the shorter length of the reads may result in worse resolution of repetitive regions. Thus, we continued only with MiSeq for short-read data for further analysis.

Compared to the short-read assemblies, the long-read assemblies had a higher overall assembly score but a lower sequence agreement score (Figure 2). Nearly 99% of long-read assemblies contained a single contig, likely due, at least in part, to the length of the reads generated and the absence of any fragmentation steps. However, the long-read assemblies were not consistent across runs, with only 30% of samples having a sequence agreement score of 1. Most of the samples with sequence agreement scores of 0 appeared to vary only by small (≤5 bp) insertions and deletions (collectively, indels), but in a few cases, there appeared to be a multiplicity issue where the length of one assembly was 2-3x longer than the assemblies from other runs (Plasmids 3121 and 3132). If a researcher’s goal is only a high-level structural confirmation of the plasmid (i.e., was my gene of interest inserted?), then such assemblies may be sufficient. However, the inconsistency of the runs is a matter of concern both in terms of overall trust in the accuracy of long-read assemblies and when detailed sequence verification is required.

The hybrid assembly pipeline produced the best assemblies overall. The overall assembly score (81%) was slightly lower than the long-read assemblies, due to the three failed short-read library preparations and the presence of more fragmented assemblies. This similarity to the short-read assemblies is consistent with the hybrid approach relying on the short-reads to establish an initial scaffold onto which the long-reads are assembled. Nevertheless, nearly 87% of samples had a good sequence agreement score, significantly higher than both short-read and long-read assemblies, highlighting the ability of the hybrid approach to reproducibly generate high quality assemblies that leverage the strengths of both technologies. There were only 3 samples that received a sequence agreement score of 0, and these had not been resolved by short-read or long-read methods either (Plasmids 3127, 3130, and 3132).

Of note, short-read assembly of the two-plasmid mixture (Plasmid 3589) resulted in either a singular plasmid of about 6600 bp or the assembly contained three contigs, seemingly representing a backbone and two inserts (Figure 2A). Several of the long-read and hybrid assemblies returned plasmids around 7400 bp and 6700 bp. Each method seemed to struggle with resolving two highly similar structures.

Given that the long-read assemblies were less robust than short-read and hybrid assemblies, we compared the results we obtained from PlasCAT to assemblies generated from Epi2ME, a long-read *de novo* assembly tool recommended by Oxford Nanopore. Epi2ME failed to produce an assembly in two cases (Supplementary Table S3), while PlasCAT always returned an assembly, albeit sometimes fragmented. Aside from these runs, 97% assemblies were successful; however, it appears that Epi2ME is restricted to returning only a single contig, which could have inflated this score. With almost 55% of samples having sequence assembly scores of 1, Epi2ME performed slightly better than PlasCAT for long-read assemblies but did not perform as well as the short-read or hybrid assemblies.

Both PlasCAT and Epi2ME subset the long-read data to a particular coverage level which should improve assembly results [22, 43, 52, 57-59]. Indeed, running the PlasCAT pipeline on the first MinION run without subsampling the data did not produce any successful assemblies; they either failed or were highly fragmented, consisting of anywhere from 3 to 80 different contigs (Supplementary Table S4). The one exception was the two-plasmid mixture (Plasmid 3589) which produced two reasonable contigs sized 7447 bp and 6532 bp. To subset the data, an estimated size for the plasmid must be provided to establish the number of reads needed to achieve a particular coverage level, presenting a potential barrier to *de novo* assembly. However, the default size parameters of PlasCAT and Epi2ME seemed to work well for samples of all sizes. While PlasCAT takes a single subset of the data to produce an assembly, Epi2ME uses Trycycler to generate a consensus assembly from 3 assemblies generated from 3 separate subsample sets, which likely improved its sequence agreement scores. Interestingly, re-analysis of the same data with the same parameters on Epi2ME occasionally returned different assemblies, suggesting a truly random seed used for subset generation. Although Trycycler was likely meant to accommodate this randomness, it was not intended to be a fully automated platform and will fail to produce a consensus if any assemblies are too different from one another [25]. On the other hand, PlasCAT’s long-read analysis returned the same assembly each time, suggesting a systematic randomness in how data is subset, and an increased ability to reliably reproduce assemblies.

Subsetting the data also led to more practical return times, cutting down the average time per PlasCAT assembly from about 30 minutes to 2 minutes. All samples from the same PlasCAT run are executed in parallel, resulting in a fast turnaround time for processing large datasets. On the other hand, each Epi2ME assembly took a few minutes per sample, and they are run in succession, so each set of 24 assemblies took two to three hours to complete. Within PlasCAT, long-read assembly was the fastest at only about 2 minutes per assembly, while each short-read assembly took, on average, 11 minutes to complete, ranging from 90 seconds to nearly 25 minutes. The hybrid assemblies took significantly longer: at least five minutes and up to 48 minutes.

### *De Novo* Assembly Reveals Deviations from Reference Sequence

Some plasmids were easily assembled by any method and matched exactly to the reference sequence (Figure 3A). However, short-read sequencers are known to have biases associated with highly repetitive and/or GC-rich regions [32, 38], thus we expected that some short-read assemblies may not be representative of the sample. Indeed, aligning the assemblies generated from the short-reads to the reference sequence revealed significant gaps in some assembly sequences (Figure 3B). While long-read sequencers are better able to resolve repetitive and GC-rich regions, they have historically been marred by high error rates, can introduce indels, and appear to depend greatly on how the data is processed (i.e., subsetting, choice of tool) [47, 60]. Thus, hybrid assemblies are expected to resolve the gaps seen in short-read assemblies by using long-read data, while also leveraging the high accuracy of short reads to prevent indels and mismatches.

**Figure 3:**
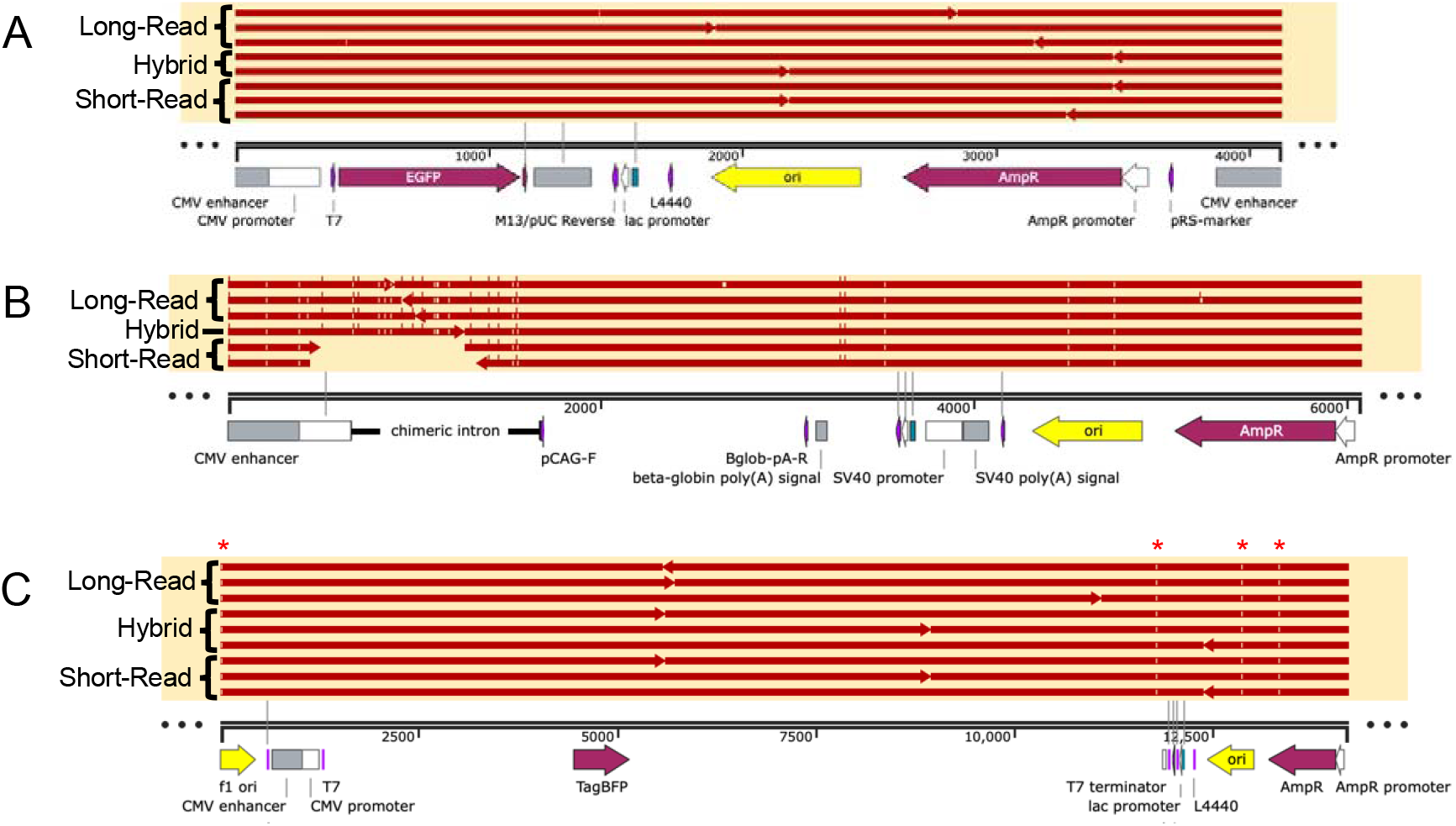
Alignment of assembly sequences to a reference sequence. **(A)** Representative example of a plasmid that is correctly assembled by any method (Plasmid 3125). **(B)** Short-read assemblies consistently miss GC-rich regions that are resolved by long-read or hybrid assemblies (Plasmid 3131). **(C)** Some deviations between assembly sequences and the reference sequence are consistent across methods (Plasmid 3133), indicating true mutations (red asterisks). Only successful assemblies were aligned to the reference sequence (1-3 per method). Each red line represents 1 assembly. Insertions, relative to the reference, appear as lines above the red line, while deletions and mismatches appear as gaps in the red line.

We found that all assembly methods resulted in some assemblies that had indels or mismatches compared to the reference sequence (Supplementary Table S5, Table 1). The short-read assemblies, and therefore the hybrid assemblies, were more likely to fail or be fragmented than the long-read assemblies. The short-read assemblies were also the most likely to have large indels, typically entire fragments missing (Table 1). However, long-read assemblies had more small indels (≤ 5 bp) and mismatches than the other methods. There were 10 plasmids where at least one long-read assembly had a deviation from the reference sequence, although the short-read and hybrid assemblies matched the reference perfectly. In all cases except one, these assemblies contained indels, mostly small (Supplementary Table S5). There were also two samples that had assemblies that matched the reference length exactly but had several mismatches, which we did not encounter in the short-read or hybrid assemblies.

**Table 1.**
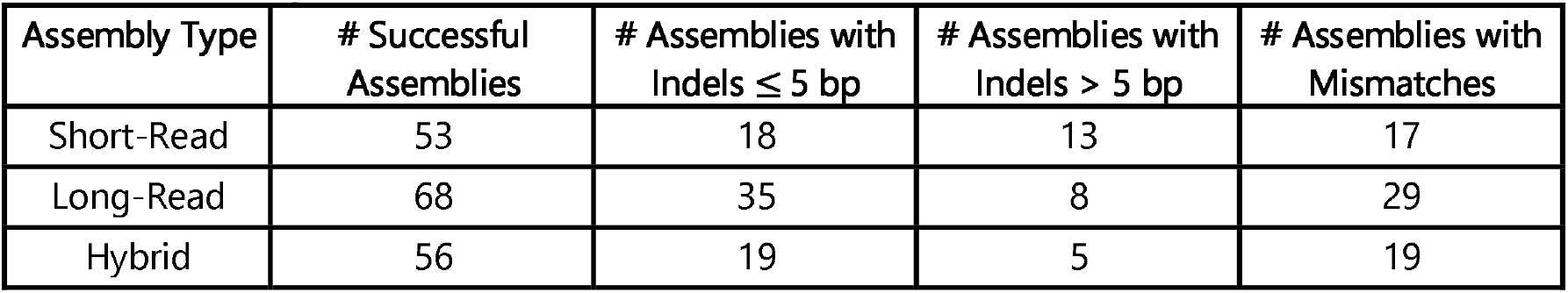
Summary of errors in plasmid assemblies compared to the reference sequence.

| Assembly Type | # Successful Assemblies | # Assemblies with Indels $\leq 5$ bp | # Assemblies with Indels $> 5$ bp | # Assemblies with Mismatches |
| --- | --- | --- | --- | --- |
| Short-Read | 53 | 18 | 13 | 17 |
| Long-Read | 68 | 35 | 8 | 29 |
| Hybrid | 56 | 19 | 5 | 19 |

There were 9 plasmids that had deviations from the reference sequence in all three types of assemblies (Supplementary Table S5). Of these, 5 plasmids consistently deviated from the reference sequence (Figure 3C). For example, the reference for Plasmid 3133 was 14194 bp, but all 9 assemblies had 6 identical mismatches and 2 small deletions resulting in a 14191 bp plasmid. In addition, 8 of the 9 assemblies for Plasmid 3121 showed a 19 bp deletion corresponding to the T7 promoter. For all 5 samples, any successful assemblies that did not exactly match the others were long-read assemblies that supported the deviations but had additional indels.

In the 4 other cases, there were deviations from the reference sequence, but it was not clear what the true sequence was. For example, Plasmid 3131 usually returned a 6077 or 6078 bp from long-read or hybrid assemblies; some errors were consistent, resulting in a plasmid roughly 13 bp larger than the reference, but a consensus could not be reached (Supplementary Table S5). All 4 plasmids showed large (>750 bp) gaps in the short-read assemblies corresponding to the chicken β-actin (CAG) promoter [61] and adjacent chimeric intron (Figure 3B, Supplementary Table S5). This region is highly repetitive and has a GC content of 73%. This region also contained the discrepancies that could not be resolved between the long-read and hybrid assemblies.

Sanger sequencing primers were designed to target a few regions containing discrepancies (Supplementary Table S1). The Sanger reactions allowed us to confirm several deviations from the reference sequence (Figure 4). For example, it confirmed a 1 bp deletion near the chicken β-actin promoter in Plasmid 3132 (Figure 4A). It also confirmed a 2 bp insertion in a section of repeated Gs in the same promoter in Plasmid 3127, which was found in one hybrid assembly; the other assemblies had a variable number of Gs and the short-read assemblies missed the region completely (Figure 4B). Given the high level of support across methods for certain discrepancies along with our Sanger data, we excluded what seemed to be true deviations, recalculated the number of assemblies with errors, and found that hybrid assemblies had the fewest remaining errors of all types (Table 2).

**Table 2.**
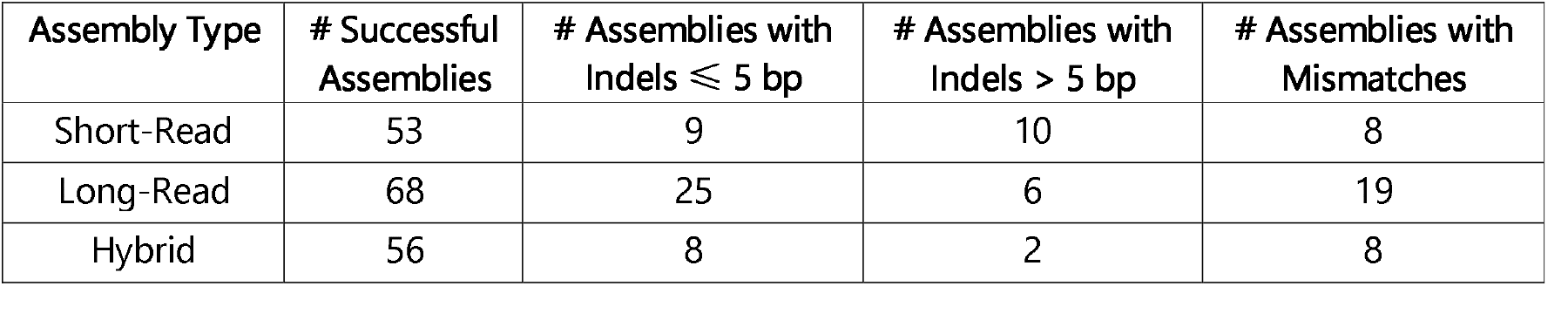
Summary of errors in plasmid assemblies compared to the reference sequence, excluding true deviations.

| Assembly Type | # Successful Assemblies | # Assemblies with Indels ≤ 5 bp | # Assemblies with Indels > 5 bp | # Assemblies with Mismatches |
| --- | --- | --- | --- | --- |
| Short-Read | 53 | 9 | 10 | 8 |
| Long-Read | 68 | 25 | 6 | 19 |
| Hybrid | 56 | 8 | 2 | 8 |

**Figure 4:**
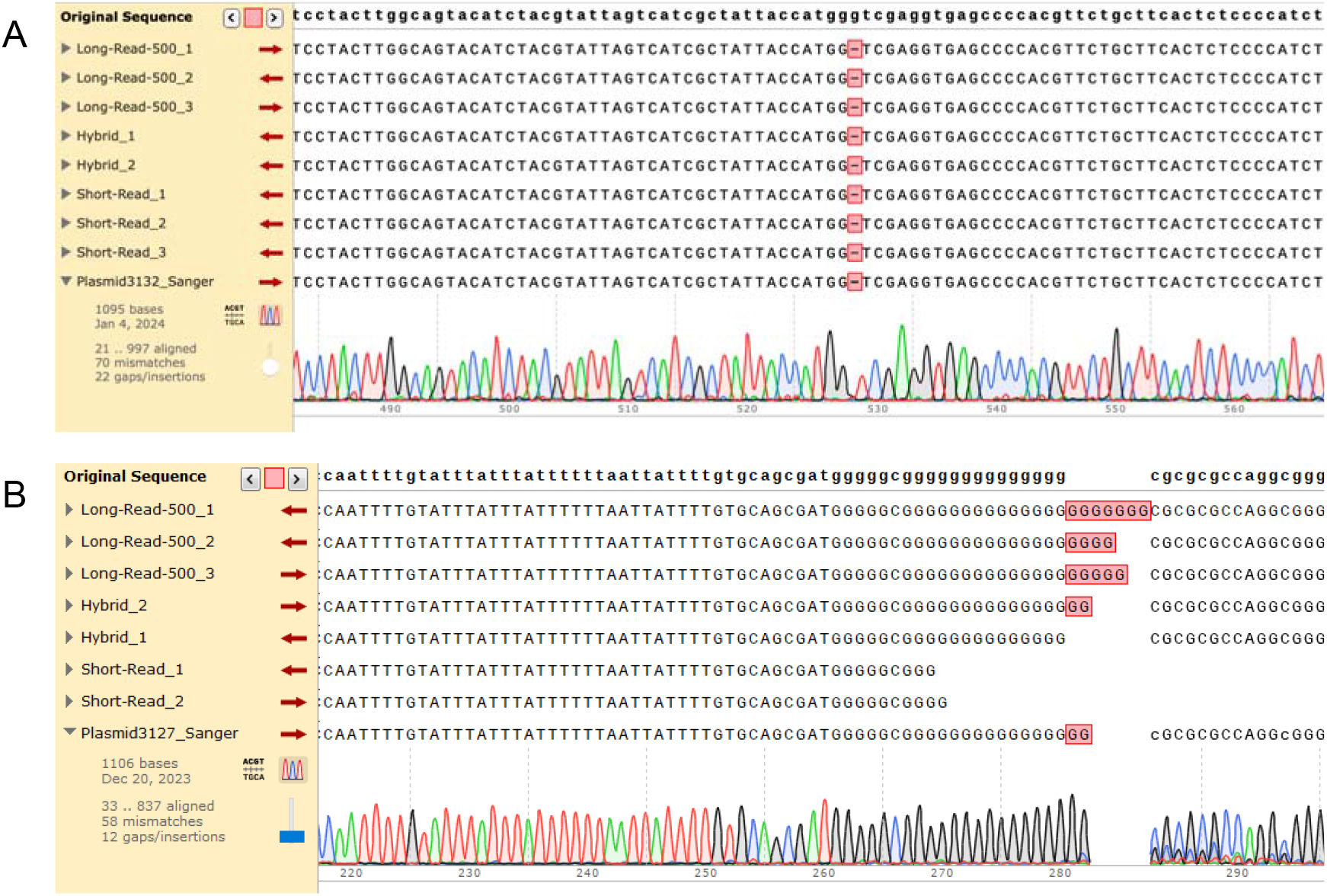
Sanger confirmation of discrepancies compared to reference sequence. All successful assemblies were aligned to the reference sequence. (A) Sanger sequencing confirmed a 1 bp deletion in the sequence of Plasmid 3132 that was found in all assemblies. (B) Sanger sequencing confirmed a 2 bp insertion in the sequence of Plasmid 3127, which was found in only one hybrid assembly. The 2 short-read assemblies missed this region entirely.

One may assume that if a reference sequence is available, performing a reference-based assembly would lead to improved results over a *de novo* assembly. We used MIRA [36, 62-64] to perform reference-based assembly of short reads for the 9 plasmids where the *de novo* assemblies deviated from the reference. A detailed discussion of these results is provided in the supplementary data (Supplementary Discussion). Briefly, the *de novo* assemblies were much closer to the expected size for all samples (Supplementary Table S6). Some of the deviations found in *de novo* assemblies were supported by the reference-based assemblies, however, there were several cases where the reference-based assembly showed evidence of reference bias: the assembly matched the reference sequence even when Sanger data supported the deviations found in the *de novo* assemblies (Supplementary Figure S2). In addition, reference-based assemblies frequently contained non-standard nucleotides, even when Sanger data showed clean peaks. These findings suggest that *de novo* assemblies are more accurate than reference-based assemblies, which is especially powerful since accurate reference sequences may not always be available.

## DISCUSSION

Calls for consistent, accurate sequence verification have been left unanswered for too long [11, 13]. Even when researchers may not expect single base pair accuracy to be important, it is important to remember that even single base pair changes can have unintended biological effects, exemplified by SNPs, various diseases, and the sequence similarity between certain fluorescent proteins [65, 66]. We evaluated several approaches for plasmid assembly in terms of their ability to produce a successful (single contig) assembly as well as to reproducibly assemble a plasmid across three technical replicates of a 24-plasmid library. In several cases, the *de novo* assemblies for a plasmid consistently differed from the reference sequence, regardless of sequencing method, and several of these deviations were confirmed by Sanger sequencing. We found *de novo* assemblies to be more accurate than reference-based assemblies. Providing a reference sequence sometimes led to reference bias wherein the reference-based assembly preferentially matched the reference sequence, even when Sanger sequencing confirmed a true deviation. Amongst *de novo* assemblies, the hybrid approach, leveraging the high accuracy of short reads with the ability of long reads to resolve complex regions, led to the best, most reproducible assemblies.

Although Sanger sequencing continues to be the gold standard for accurately resolving known short sequences [67], it requires a reference sequence in order to generate primers and increasing the number of reactions to cover larger sections becomes costly and time-consuming (Supplementary Table S7). Thus, it is not a practical approach for *de novo* plasmid assembly. Long-read plasmid assembly is best-suited for obtaining a complete structural overview of a plasmid, but issues like indels may make assemblies less trustworthy or mean that small alterations may be missed in the absence of additional replicates. Short-read plasmid assembly may be sufficient for plasmids that are not highly repetitive and with an overall and parts-level GC-content around 50%, otherwise large fragments may be missing or the assemblies may be fragmented. Although short-read assemblies can outperform long-read assemblies, it is worth noting that short-read sequencing on the Illumina MiSeq is more expensive per sample than long-read sequencing on the ONT MinION (Supplementary Table S7). While one may wish to argue that short– or long-read sequencing on their own can provide sufficient sequence verification, the hybrid approach not only leverages the high accuracy of short-reads, but the ability of long-reads to resolve complex regions. The cost associated with this choice will be the highest, but in return, reproducibility and confidence will skyrocket. When deviations from the expected sequence arise during hybrid assembly, confidence can be gained by sequencing additional independent replicates of your plasmid. Given the propensity of DNA to mutate, errors could arise in one bacterial colony and not another, and frequent sequencing can help detect which mutations arose and when. Sanger sequencing can provide additional confirmation of potential errors.

However, it should be emphasized that certain highly repetitive, GC-rich sequences remained difficult to resolve to single base pair accuracy by any of the bioinformatics pipelines, with the short-read data performing the worst in these areas. Specifically, there were 4 plasmids containing a GC-rich region encompassing a CAG promoter and adjacent chimeric intron, which could not be resolved by any assembly method. Although targeted Sanger sequencing may resolve these sequences with good confidence (Figure 4), regions with GC-content outside the typical (40-60%) range as well as repetitive regions that can form hairpins can present challenges for Sanger, short-read, and long-read sequencing alike [2, 41, 67]. If some genetic parts are unreliable to sequence regardless of the method, especially to single base pair accuracy, these parts must be cataloged and flagged as difficult sequences. Ideally, these parts would be replaced with others that can perform the same function but are more readily sequenced.

This work was performed with the large-scale production and verification of high-throughput plasmid libraries in mind. With increasing ease and availability of DNA synthesis and sequencing technologies [68], there is increasing demand for high-throughput production methods to produce hundreds to thousands of plasmids. The major bottleneck in the production of these libraries is the sequence verification step [2, 10, 13]. Tools such as Trycycler that require manual intervention for users to reliably generate good assemblies may be useful for difficult-to-assemble plasmids but become impractical to scale up [3, 69]. Both PlasCAT and Epi2ME are into easy-to-use full-service workflows suitable for high-throughput *de novo* plasmid sequence verification. Given that a hybrid approach is superior for *de novo* plasmid assembly, it is critical that assembly tools can accommodate data from multiple sources, making a vendor-independent solution like PlasCAT appealing.

The pipelines described in this manuscript address a critical need for better plasmid library validation, generating reliable data faster, and they make it easier for non-technical users to carry out complex bioinformatics analyses. As sequencing technologies and bioinformatics tools continue to improve, further work is needed to optimize analysis pipelines for *de novo* plasmid assembly as well as improve verification and documentation practices surrounding plasmids. Recent work has led to the development of DNA signatures that can embed identifying information directly into plasmid sequences to facilitate simpler plasmid verification using only *de novo* assembly [1, 70]. By incorporating a compressed version of a sequence into a signature and inserting this into a plasmid, plasmids can be instantly verified against the original reference sequence even with no prior knowledge of the sequence, emphasizing the need for and power of *de novo* assembly [1, 70].

## DATA AVAILABILITY

All data generated for this manuscript are publicly available in a repository and can be accessed at https://figshare.com/s/cb61b237859049e68e52.

Supplementary Information and Data are available at NAR online.

## AUTHOR CONTRIBUTIONS

Sarah Hernandez: Conceptualization, Methodology, Formal Analysis, Visualization, Writing— original draft & editing. Casey-Tyler Berezin: Conceptualization, Formal analysis, Software, Writing—review & editing. Katie Miller: Formal analysis, Writing—review. Samuel Peccoud: Software. Jean Peccoud: Conceptualization, Methodology, Writing—review & editing.

## ACKNOWLEDGEMENTS

Figure 1 was created with BioRender.com.

## REFERENCES

1. Gallegos, J.E., et al., Securing the Exchange of Synthetic Genetic Constructs Using Digital Signatures. ACS Synth. Biol., 2020. 9(10): p. 2656–2664.

2. Gallegos, J.E., et al., Rapid, robust plasmid verification by de novo assembly of short sequencing reads. Nucleic Acids Research, 2020. 48(18).

3. Brown, S.D., et al., Complete sequence verifcation of plasmid DNA using the Oxford Nanopore Technologies’ MinION device. BMC Bioinformatics, 2023. 24(116).

4. Rozwandowicz, M., et al., Plasmids carrying antimicrobial resistance genes in Enterobacteriaceae. Journal of Antimicrobial Chemotherapy, 2018. 73: p. 1121–1137.

5. Cameron, D.E., C.J. Bashor, and J.J. Collins, A brief history of synthetic biology. Nature Reviews Microbiology, 2014. 12(5): p. 381–390.

6. Munnelly, K., Engineering for the 21st Century: Synthetic Biology. ACS Synth. Biol., 2013(2): p. 213–215.

7. Peccoud, J., Synthetic Biology: fostering the cyber-biological revolution. Synthetic Biology, 2016. 1(1).

8. Ghaffarifar, F., Plasmid DNA vaccines: where are we now. Drugs Today, 2018. 54(5): p. 315–33.

9. Shapland, E.B., V. Holmes, and C.D. Reeves, Low-Cost, High-Throughput Sequencing of DNA Assemblies Using a Highly Multiplexed Nextera Process. ACS Synthetic Biology, 2015. 4(7): p. 860–866.

10. Peccoud, J., Data sharing policies: share well and you shall be rewarded. Synthetic Biology, 2021. 6(1): p. ysab028.

11. Peccoud, J., et al., Essential information for synthetic DNA sequences. Nature Biotechnology, 2011. 29(1): p. 22–22.

12. Peccoud, J., et al., Cyberbiosecurity: from naive trust to risk awareness. Trends in biotechnology, 2018. 36(1): p. 4–7.

13. Thuronyi, B.W., E.A. DeBenedictis, and J.E. Barrick, No assembly required: Time for stronger, simpler publishing standards for DNA sequences. Plos Biology, 2023. 21(11): p. e3002376.

14. Antipov, D., et al., plasmidSPAdes: assembling plasmids from whole genome sequencing data. Bioinformatics, 2016. 32(22): p. 3380–3387.

15. Rozov, R., et al., Recycler: an algorithm for detecting plasmids from de novo assembly graphs. Bioinformatics, 2017. 33(4): p. 475–482.

16. Antipov, D., et al., Plasmid detection and assembly in genomic and metagenomic data sets. Genome Res, 2019. 29(6): p. 961–968.

17. Gomi, R., K.L. Wyres, and K.E. Holt, Detection of plasmid contigs in draft genome assemblies using customized Kraken databases. Microb Genom, 2021. 7(4).

18. Pellow, D., et al., SCAPP: an algorithm for improved plasmid assembly in metagenomes. Microbiome, 2021. 9(1): p. 144.

19. Gupta, S.K., S. Raza, and T. Unno, Comparison of de-novo assembly tools for plasmid metagenome analysis. Genes & Genomics, 2019. 41(9): p. 1077–1083.

20. Wick, R.R., et al., Unicycler: resolving bacterial genome assemblies from short and long sequencing reads. PLoS computational biology, 2017. 13(6): p. e1005595.

21. Tang, X., et al., PLASMe: a tool to identify PLASMid contigs from short-read assemblies using transformer. Nucleic Acids Research, 2023. 51(15): p. e83–e83.

22. Bouras, G., et al., Plassembler: an automated bacterial plasmid assembly tool. Bioinformatics, 2023. 39(7).

23. Berbers, B., et al., Combining short and long read sequencing to characterize antimicrobial resistance genes on plasmids applied to an unauthorized genetically modifed Bacillus. Nautre Research, 2020. 10.

24. Johnson, J., M. Soehnlen, and H.M. Blankenship, Long read genome assemblers struggle with small plasmids. Microbial Genomics, 2023. 9(5).

25. Wick, R.R., et al., Trycycler: consensus long-read assemblies for bacterial genomes. Genome Biology, 2021. 22.

26. Chen, N.-C., et al., Reference flow: reducing reference bias using multiple population genomes. Genome Biology, 2021. 22(1).

27. Valiente-Mullor, C., et al., One is not enough: On the effects of reference genome for the mapping and subsequent analyses of short-reads. PLOS Computational Biology, 2021. 17(1): p. e1008678.

28. Lau, J. Reference bias: Challenges and solutions. SevenBridges Blog, 2017.

29. Berezin, C.-T., et al. PlasCAT: Plasmid Cloud Assembly Tool. 2023.

30. Epi2ME Labs. Epi2ME Labs Blog. 2023 12/06/2023.

31. Peccoud, S., et al., PlasCAT: Plasmid Cloud Assembly Tool. Bioinformatics, 2024.

32. Mardis, E.R., Next-Generation Sequencing Platforms. Annu. Rev. Anal. Chem., 2013. 6: p. 287–303.

33. Hu, T., et al., Next-generation sequencing technologies: An overview. Human Immunology, 2021. 82(11): p. 801–811.

34. Heather, J.M. and B. Chain, The sequence of sequencers: The history of sequencing DNA. Genomics, 2016. 107(1): p. 1–8.

35. Peccoud, J., et al., Targeted development of registries of biological parts. Plos one, 2008. 3(7): p. e2671.

36. Wilson, M.L., et al., Sequence verification of synthetic DNA by assembly of sequencing reads. Nucleic Acids Research, 2012. 41(1): p. e25–e25.

37. Buermans, H.P.J. and J.T.d. Dunnen, Next generation sequencing technology: Advances and applications. Biochimica et Biophysica Acta, 2014. 1842(10): p. 1932–1941.

38. Tilak, M.-K., et al., Illumina Library Preparation for Sequencing the GC-Rich Fraction of Heterogeneous Genomic DNA. Genome Biology and Evolution, 2018. 10(2): p. 616–622.

39. Liao, X., et al., Current challenges and solutions of de novo assembly. Quantitative Biology, 2019. 7: p. 90–109.

40. Aird, D., et al., Analyzing and minimizing PCR amplification bias in Illumina sequencing libraries. Genome Biology, 2011. 12.

41. Browne, P.D., et al., GC bias affects genomic and metagenomic reconstructions, underrepresenting GC-poor organisms. GigaScience, 2020. 9(2).

42. Zhao, W., et al., Oxford nanopore long-read sequencing enables the generation of complete bacterial and plasmid genomes without short-read sequencing. Front. Microbiol. Sec. Evolutionary and Genomic Microbiology, 2023. 14.

43. De Maio, N., et al., Comparison of long-read sequencing technologies in the hybrid assembly of complex bacterial genomes. Microbial genomics, 2019. 5(9): p. e000294.

44. Xia, Y., et al., Strategies and tools in illumina and nanopore‐integrated metagenomic analysis of microbiome data. iMeta, 2023. 2(1).

45. Khrenova, M.G., et al., Nanopore sequencing for de novo bacterial genome assembly and search for single-nucleotide polymorphism. International Journal of Molecular Sciences, 2022. 23(15): p. 8569.

46. Gallegos, J.E., et al., Challenges and opportunities for strain verification by whole-genome sequencing. Scientific Reports, 2020. 10(1): p. 5873.

47. Amarasinghe, S.L., et al., Opportunities and challenges in long-read sequencing data analysis. Genome Biology, 2020. 21(30).

48. ThermoFisher Scientific, Expi293™ Expression System USER GUIDE. 2020, ThermoFisher Scientific. p. 1–32.

49. Janeway Ca Jr, T.P., Walport M, et al., The structure of a typical antibody molecule, in Immunobiology: The Immune System in Health and Disease. 2001, New York: Garland Science.

50. Bolger, A.M., M. Lohse, and B. Usadel, Trimmomatic: a flexible trimmer for Illumina sequence data. Bioinformatics, 2014. 30(15): p. 2114–2120.

51. Hall, M.B., Rasusa: Randomly subsample sequencing reads to a specified coverage. Journal of Open Source Software, 2022. 7(69): p. 3941.

52. Lonardi, S., et al., When less is more: ‘slicing’ sequencing data improves read decoding accuracy and de novo assembly quality. Bioinformatics, 2015. 31(18): p. 2972–2980.

53. Wick, R.R. and P. Menzel. Filtlong. Filtlong 2021 [cited 2024.

54. Vaser, R., et al., Fast and accurate de novo genome assembly from long uncorrected reads. Genome Res., 2017. 27(5): p. 737–746.

55. Kolmogorov, M., et al., Assembly of long, error-prone reads using repeat graphs. Nature Biotechnology, 2019. 37: p. 540–546.

56. Wright, C., et al. epi2me-labs/wf-denovo-assembly. epi2me-labs 2022 11/16/2023 [cited 2023; Available from: https://github.com/epi2me-labs/wf-denovo-assembly.

57. Murigneux, V., et al., Comparison of long-read methods for sequencing and assembly of a plant genome. GigaScience, 2020. 9(12).

58. Wick, R.R. and K.E. Holt, Benchmarking of long-read assemblers for prokaryote whole genome sequencing. F1000Research, 2021. 8: p. 2138.

59. Hall, M.B., Rasusa: Randomly subsample sequencing reads to a specified coverage. Journal of Open Source Software, 2022. 7(69).

60. Boostrom, I., et al., Comparing long-read assemblers to explore the potential of a sustainable low-cost, low-infrastructure approach to sequence antimicrobial resistant bacteria with oxford nanopore sequencing. Frontiers in Microbiology, 2022. 13: p. 796465.

61. Alexopoulou, A.N., J.R. Couchman, and J.R. Whiteford, The CMV early enhancer/chicken β actin (CAG) promoter can be used to drive transgene expression during the differentiation of murine embryonic stem cells into vascular progenitors. BMC Cell Biology, 2008. 9(1): p. 2.

62. Chevreux, B., T. Wetter, and S. Suhai. Genome Sequence Assembly Using Trace Signals and Additional Sequence Information. in German Conference on Bioinformatics. 1999.

63. Cock, P.J., et al., Galaxy tools and workflows for sequence analysis with applications in molecular plant pathology. PeerJ, 2013. 1: p. e167.

64. The Galaxy Community, The Galaxy platform for accessible, reproducible and collaborative biomedical analyses: 2022 update. Nucleic Acids Research, 2022. 50(W1): p. W345–W351.

65. Morise, H., et al., Intermolecular Energy Transfer in the Bioluminescent System of Aequorea. Biochemistry, 1974. 13(12): p. 2656–2662.

66. Weiner, M.P. and T.J. Hudson, Introduction to SNPs: discovery of markers for disease. Biotechniques, 2002. 32(Sup): p. S4–S13.

67. Crossley, B.M., et al., Guidelines for Sanger sequencing and molecular assay monitoring. 2020. 32(6): p. 767–775.

68. Hughes, R.A. and A.D. Ellington, Synthetic DNA Synthesis and Assembly: Putting the Synthetic in Synthetic Biology. Cold Spring Harbor Perspectives in Biology, 2017. 9(1).

69. Wick, R.R., L.M. Judd, and K.E. Holt, Assembling the perfect bacterial genome using Oxford Nanopore and Illumina sequencing. PLOS Computational Biology, 2023. 19(3): p. e1010905.

70. Berezin, C.-T., et al., Cryptographic approaches to authenticating synthetic DNA sequences. Trends in Biotechnology, 2024.

