## Supplementary Information for "Sequencing Strategy to Ensure Accurate Plasmid Assembly"

**Supplementary Information and Data**

Supplementary Discussion of Reference-Based Assembly

Supplementary Tables S1-S7

Supplementary Figures S1-S2

Supplementary References

_____________________________________________________________________________________________

**Supplementary Discussion of Reference-Based Assembly**

We considered the possibility that a reference-based assembly approach may produce better results than *de novo* assembly approach. Interestingly, there are few tools built for this purpose as most recently developed assembly tools rely on *de novo* methods. We used MIRA v3.4 *via* the public NanoGalaxy server at nanopore.usegalaxy.eu to perform reference-based assembly of short reads (1-4). The assemblies were generated by using “Mapping” mode, “Accurate” grade, and providing the reference sequence. Short reads from the first run were used in all cases and Sanger data was additionally used where available. We focused only on the 9 plasmids where the *de novo* assemblies deviated from the reference.

All the reference-based assemblies were, on average, 700 bp longer than the expected reference sequences, lending initial support for the *de novo* approach which resulted in assemblies closer to the expected size (Supplementary Table S6). In addition, the reference-based assemblies often included non-standard nucleotide abbreviations, e.g., R representing a purine (A or G) and M representing A or C. While these could represent a mixed population of sequences, the Sanger data did not support a mixed population as there were clear peaks supporting those nucleotides (Supplementary Figure S2A). Thus, it is more likely that the reference-based assembly simply could not resolve all sequences.

In some cases, the reference-based assemblies supported the deviations we found from the reference in the *de novo* assemblies. For example, all the deviations found in Plasmid 3123 were supported by the reference-based assembly. When the two assembly types did not agree, the reference-based assembly either matched the reference sequence exactly or contained a non-standard nucleotide (Supplementary Figure S2A, S2B). Although the reference-based short-read assemblies did not contain the same large gaps as the *de novo* short-read assemblies, this was most likely due to the use of the reference sequence to supplement the assembly rather than the ability to assemble the reads in that region.

Sanger reads can be provided alongside short reads to generate the reference-based assemblies. In some cases, providing the Sanger data improved the resulting MIRA assembly such that it matched the Sanger data as well as the *de novo* assemblies (Supplementary Figure S2B). However, in other cases, the reference-based assembly seemed to prioritize a match to the reference sequence and completely ignored the provided Sanger data (Supplementary Figure S2C). Given that Sanger sequencing is still considered the gold standard for resolving short sequences with high accuracy, the fact that the reference-based assemblies often did not match the Sanger data suggests that reference-based assemblies are not as accurate as *de novo* assemblies.

_____________________________________________________________________________________________

**Supplementary Tables**

**Table S1. Primers used for Sanger sequencing.**

| **Plasmid** | **Sequence** | **Reverse Compliment** | **Size (bp)** | **Melting Temperature (°C)** | **GC%** |
| --- | --- | --- | --- | --- | --- |
| 3127 | ACGGTAAACTGCCCACTTGGC | GCCAAGTGGGCAGTTTACCGT | 21 | 69.4 | 57.1 |
| 3127 | GCGCTAATTACAGCCCGGAGG | CCTCCGGGCTGTAATTAGCGC | 21 | 70.6 | 61.9 |
| 3133 | GCCCTGGACGGCGTACTTCC | GGAAGTACGCCGTCCAGGGC | 20 | 70 | 70 |
| 3133 | GGCTGTCGGGCATTCCTTGC | GCAAGGAATGCCCGACAGCC | 20 | 73.3 | 65 |
| 3131 | CTGGCTGACCGCCCAACGAC | GTCGTTGGGCGGTCAGCCAG | 20 | 74.7 | 70 |
| 3131 | CCACGCACACACACACGCAC | GTGCGTGTGTGTGTGCGTGG | 20 | 73.1 | 65 |
| 3130 | GGGCGACACGGAAATGTTGA | TCAACATTTCCGTGTCGCCC | 20 | 71.1 | 65 |
| 3130 | GCCCGCTCACCTGTGGGAGT | ACTCCCACAGGTGAGCGGGC | 20 | 73.4 | 70 |

**Table S2. Summary of results from iSeq assemblies.**

| **Plasmid #** | **iSeq Assembly Notes** |
| --- | --- |
| 3125 | Assembly contains 1 contig of length 4119. Total time 358.069 seconds. |
| 3137 | Assembly contains 1 contig of length 4875. Total time 371.905 seconds. |
| 3139 | Assembly contains 1 contig of length 4908. Total time 218.451 seconds. |
| 3113 | Assembly contains 1 contig of length 4908. Total time 356.412 seconds. |
| 3144 | ** Assembly contains 3 fragments (4911, 1415, 1069) totalling 7395 bases. ** Warning! Fragment(s) may be linear: #2, 3 Total time 390.642 seconds. |
| 3148 | Assembly contains 1 contig of length 4947. Total time 216.045 seconds. |
| 3122 | Assembly contains 1 contig of length 4966. Total time 354.312 seconds. |
| 3118 | Assembly contains 1 contig of length 5041. Total time 207.751 seconds. |
| 3117 | Assembly contains 1 contig of length 5050. Total time 364.417 seconds. |
| 3115 | Assembly contains 1 contig of length 6815. Total time 221.434 seconds. |
| 2101 | Assembly contains 1 contig of length 6824. Total time 375.697 seconds. |
| 3123 | Assembly contains 1 contig of length 6832. Total time 356.349 seconds. |
| 3124 | Assembly contains 1 contig of length 6819. Total time 370.806 seconds. |
| 3127 | Assembly contains 1 contig of length 6652. ** Warning! Fragment(s) may be linear: #1 Total time 368.463 seconds. |
| 3121 | Assembly contains 1 contig of length 7765. Total time 354.331 seconds. |
| 3116 | ** Assembly contains 3 fragments (4454, 4026, 138) totalling 8618 bases. ** Warning! Fragment(s) may be linear: #1, 2, 3 Total time 372.764 seconds. |
| 3126 | Assembly contains 1 contig of length 4611. Total time 219.505 seconds. |
| 3131 | Assembly contains 1 contig of length 5295. ** Warning! Fragment(s) may be linear: #1 Total time 389.099 seconds. |
| 3132 | Assembly contains 1 contig of length 5467. ** Warning! Fragment(s) may be linear: #1 Total time 222.370 seconds. |
| 3130 | Assembly contains 1 contig of length 10229. ** Warning! Fragment(s) may be linear: #1 Total time 346.232 seconds. |
| 3135 | ** Assembly contains 2 fragments (10462, 3005) totalling 13467 bases. ** Warning! Fragment(s) may be linear: #1, 2 Total time 212.885 seconds. |
| 3133 | Assembly contains 1 contig of length 14191. Total time 375.158 seconds. |
| 3134 | ** Assembly contains 2 fragments (12843, 2068) totalling 14911 bases. ** Warning! Fragment(s) may be linear: #1, 2 Total time 215.594 seconds. |
| 3589 | ** Assembly contains 3 fragments (5976, 1486, 778) totalling 8240 bases.   ** Filtering with minimum quality 35 and window size 9 yields 12330 filtered read pairs. ** Warning! Fragment(s) may be linear: #1, 2, 3 Total time 408.142 seconds. |

**Table S3. Summary of results from Epi2ME assemblies.**

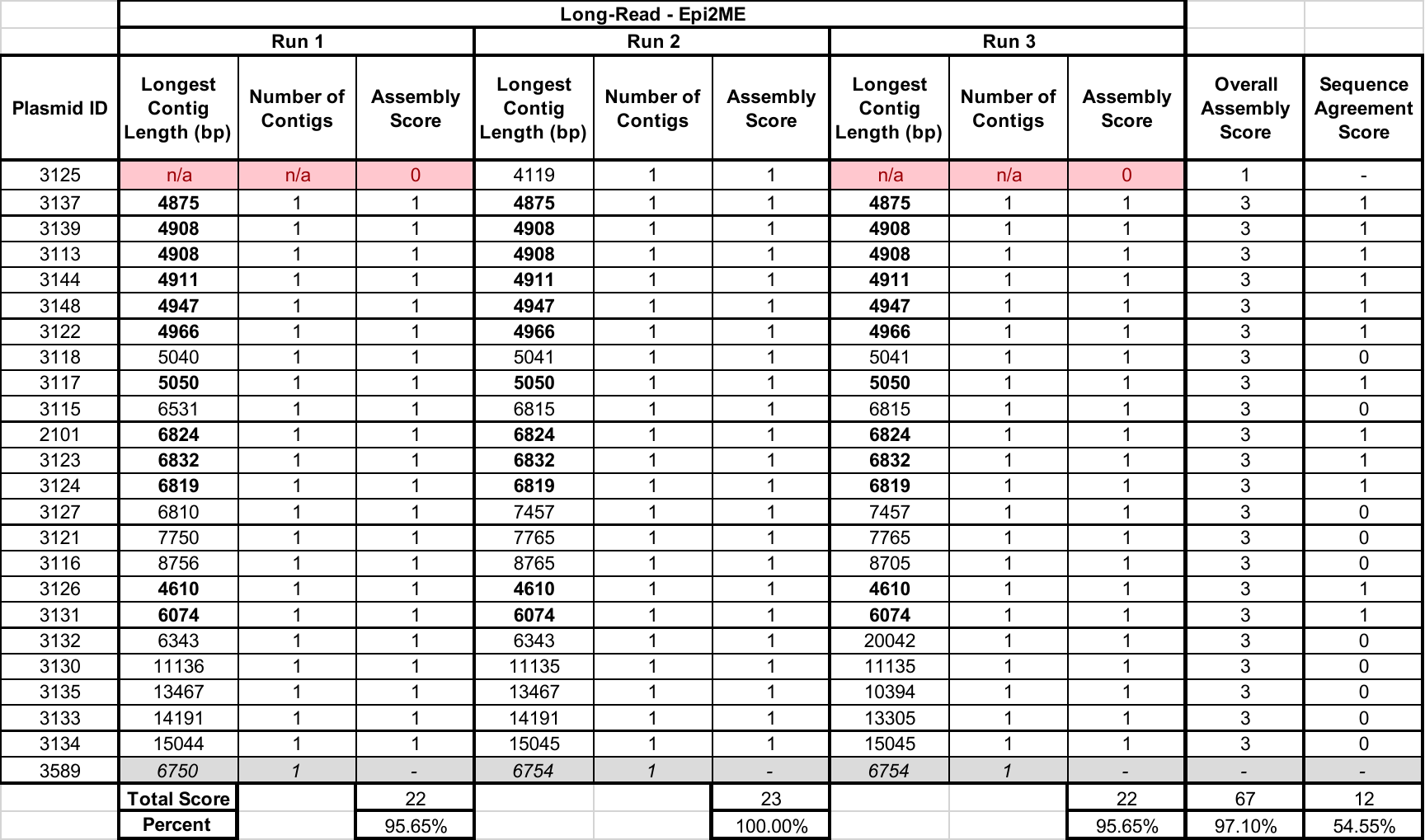

**Table S4. Summary of results of long-read assemblies without subsampling.**

| **Plasmid #** | **Number of Fragments** |
| --- | --- |
| 3125 | 63 |
| 3137 | 9 |
| 3139 | 5 |
| 3113 | 72 |
| 3144 | 54 |
| 3148 | 43 |
| 3122 | 37 |
| 3118 | 45 |
| 3117 | 50 |
| 3115 | 3 |
| 2101 | 63 |
| 3123 | 5 |
| 3124 | 80 |
| 3127 | 26 |
| 3121 | 9 |
| 3116 | 3 |
| 3126 | 36 |
| 3131 | 3 |
| 3132 | 57 |
| 3130 | Failed |
| 3135 | 70 |
| 3133 | 3 |
| 3134 | 62 |
| 3589 | 2 |

**Table S5. Summary of errors found between *de novo* assemblies and reference sequences.**

| **Plasmid ID** | **Reference Length (bp)** | **Assembly type** | **Run #** | **Assembly Length (bp)** | **Error Type (n)** | **Indel details** |
| --- | --- | --- | --- | --- | --- | --- |
| 3125 | 4119 | Long-read | 1 | 4117 | indel ≤ 5 bp (1) | -2bp |
|  |  |  | 3 | 4116 | indel ≤ 5 bp (1) | -3bp |
| 3137 | 4875 | Long-read | 1 | 4874 | indel ≤ 5 bp (1) | -1bp |
| 3139 | 4908 | Long-read | 2 | 4909 | indel ≤ 5 bp (2) mismatches (20) | +2bp & -1bp |
| 3122 | 4966 | Long-read | 2 | *4966* | mismatches (54) | - |
|  |  |  | 3 | 4965 | indel ≤ 5 bp (1) | -1bp |
| 2101 | 6824 | Long-read | 2 | 6820 | indel ≤ 5 bp (2) | -1bp & -3bp |
|  |  |  | 3 | 6823 | indel ≤ 5 bp (1) | -1bp |
| 3126 | 4611 | Long-read | 3 | 4914 | indel ≤ 5 bp (1) mismatches (2) | +3bp (rep. GCC) |
| 3115 | 6815 | Long-read | 2 | *6815* | mismatches (61) | - |
| 3137 | 4875 | Long-read | 1 | 4874 | indel ≤ 5 bp (1) | -1bp |
| 3116 | 8756 | Long-read | 1 | 8757 | indel ≤ 5 bp (1) | +1bp |
| 3135 | 13467 | Long-read | 2 | 13059 | indel ≤ 5 bp (7) indel > 5 bp (13) mismatches (5) | +535 bp, +282 bp, +92bp, +82bp, +36bp, +9bp, +3bp, +1bp, -738bp, -290bp, -224bp, -75bp, -52bp, -51bp, -8bp, -4bp, -3bp, 3 -1bp |
|  |  |  | 3 | 12067 | indel > 5 bp (2) | -808bp & -592bp |
| 3124 | 6817 | Short-read | 1 | **6819** | indel ≤ 5 bp (4) mismatches (31) | 3 +1bp (1 rep. C) & -1bp |
|  |  |  | 3 | **6819** | indel ≤ 5 bp (4) mismatches (31) | 3 +1bp (1 rep. C) & -1bp |
|  |  | Long-read | 1 | **6819** | indel ≤ 5 bp (4) mismatches (31) | 3 +1bp (1 rep. C) & -1bp |
|  |  |  | 2 | **6819** | indel ≤ 5 bp (4) mismatches (31) | 3 +1bp (1 rep. C) & -1bp |
|  |  |  | 3 | **6819** | indel ≤ 5 bp (5) mismatches (31) | 3 +1bp (1 rep. C) & -1bp |
|  |  | Hybrid | 1 | **6819** | indel ≤ 5 bp (4) mismatches (31) | 3 +1bp (1 rep. C) & -1bp |
|  |  |  | 3 | **6819** | indel ≤ 5 bp (4) mismatches (31) | 3 +1bp (1 rep. C) & -1bp |
| 3133 | 14194 | Short-read | 1 | **14191** | indel ≤ 5 bp (2) mismatches (6) | -1bp (rep. G) & -2bp (rep. G) |
|  |  |  | 2 | **14191** | indel ≤ 5 bp (2) mismatches (6) | -1bp (rep. G) & -2bp (rep. G) |
|  |  |  | 3 | **14191** | indel ≤ 5 bp (2) mismatches (6) | -1bp (rep. G) & -2bp (rep. G) |
|  |  | Long-read | 1 | **14191** | indel ≤ 5 bp (2) mismatches (6) | -1bp (rep. G) & -2bp (rep. G) |
|  |  |  | 2 | **14191** | indel ≤ 5 bp (2) mismatches (6) | -1bp (rep. G) & -2bp (rep. G) |
|  |  |  | 3 | **14191** | indel ≤ 5 bp (2) mismatches (6) | -1bp (rep. G) & -2bp (rep. G) |
|  |  | Hybrid | 1 | **14191** | indel ≤ 5 bp (2) mismatches (6) | -1bp (rep. G) & -2bp (rep. G) |
|  |  |  | 2 | **14191** | indel ≤ 5 bp (2) mismatches (6) | -1bp (rep. G) & -2bp (rep. G) |
|  |  |  | 3 | **14191** | indel ≤ 5 bp (2) mismatches (6) | -1bp (rep. G) & -2bp (rep. G) |
| 3121 | 7784 | Short-read | 1 | **7765** | indel > 5 bp (1) | -19 bp = T7 promoter |
|  |  |  | 2 | **7765** | indel > 5 bp (1) | -19 bp = T7 promoter |
|  |  |  | 3 | **7765** | indel > 5 bp (1) | -19 bp = T7 promoter |
|  |  | Long-read | 1 | **7765** | indel > 5 bp (1) | -19 bp = T7 promoter |
|  |  |  | 2 | 20878 | indel > 5 bp (2) | -19 bp = T7 promoter; multiplicity: +13113 bp |
|  |  |  | 3 | **7765** | indel > 5 bp (1) | -19 bp = T7 promoter |
|  |  | Hybrid | 1 | **7765** | indel > 5 bp (1) | -19 bp = T7 promoter |
|  |  |  | 2 | **7765** | indel > 5 bp (1) | -19 bp = T7 promoter |
|  |  |  | 3 | **7765** | indel > 5 bp (1) | -19 bp = T7 promoter |
| 3123 | 6828 | Short-read | 1 | **6832** | indel ≤ 5 bp (2) mismatches (1) | +1 bp (rep. A) & +3bp (rep. GGT) |
|  |  |  | 2 | **6832** | indel ≤ 5 bp (2) mismatches (1) | +1 bp (rep. A) & +3bp (rep. GGT) |
|  |  |  | 3 | **6832** | indel ≤ 5 bp (2) mismatches (1) | +1 bp (rep. A) & +3bp (rep. GGT) |
|  |  | Long-read | 1 | **6832** | indel ≤ 5 bp (2) mismatches (1) | +1 bp (rep. A) & +3bp (rep. GGT) |
|  |  |  | 2 | **6832** | indel ≤ 5 bp (2) mismatches (1) | +1 bp (rep. A) & +3bp (rep. GGT) |
|  |  |  | 3 | 6919 | indel ≤ 5 bp (3) indel > 5 bp (2) mismatches (5) | +1 bp (rep. A), +3bp (rep. GGT), +2bp, +23bp, +62bp |
|  |  | Hybrid | 1 | **6832** | indel ≤ 5 bp (2) mismatches (1) | +1 bp (rep. A) & +3bp (rep. GGT) |
|  |  |  | 2 | **6832** | indel ≤ 5 bp (2) mismatches (1) | +1 bp (rep. A) & +3bp (rep. GGT) |
|  |  |  | 3 | **6832** | indel ≤ 5 bp (2) mismatches (1) | +1 bp (rep. A) & +3bp (rep. GGT) |
| 3134 | 15045 | Short-read | 2 | **15044** | indel ≤ 5 bp (1) mismatches (3) | -1bp |
|  |  | Long-read | 1 | 15041 | indel ≤ 5 bp (2) mismatches (3) | -1bp & -3bp |
|  |  |  | 2 | **15044** | indel ≤ 5 bp (1) mismatches (3) | -1bp |
|  |  |  | 3 | **15044** | indel ≤ 5 bp (1) mismatches (3) | -1bp |
|  |  | Hybrid | 1 | **15044** | indel ≤ 5 bp (1) mismatches (3) | -1bp |
|  |  |  | 2 | **15044** | indel ≤ 5 bp (1) mismatches (3) | -1bp |
|  |  |  | 3 | **15044** | indel ≤ 5 bp (1) mismatches (3) | -1bp |
| 3131 | 6064 | Short-read | 1 | 5295 | indel ≤ 5 bp (8) indel > 5 bp (1) mismatches (7) | 7 +1bp, -1 bp, -775bp |
|  |  |  | 2 | 5179 | indel ≤ 5 bp (7) indel > 5 bp (1) mismatches (7) | 6 +1bp, -1bp, -890bp |
|  |  | Long-read | 1 | 6077 | indel ≤ 5 bp (15) mismatches (18) | 11 +1bp, +4bp (rep. G), +2bp, 4 -1bp |
|  |  |  | 2 | 6080 | indel ≤ 5 bp (17) indel > 5 bp (2) mismatches (14) | +14bp, -12 bp, +5bp (rep. G), +2bp, 11 +1bp, 4 -1bp |
|  |  |  | 3 | 6077 | indel ≤ 5 bp (17) mismatches (14) | 11 +1bp, +4bp (rep. G), +2bp, 4 -1bp |
|  |  | Hybrid | 1 | 6078 | indel ≤ 5 bp (16) mismatches (14) | 11 +1bp, +2bp, +4bp (rep. G), 3 -1bp |
| 3130 | 11125 | Short-read | 1 | 10229 | indel ≤ 5 bp (7) indel > 5 bp (1) mismatches (7) | 6 +1bp, -1bp, -901bp |
|  |  |  | 2 | 10270 | indel ≤ 5 bp (7) indel > 5 bp (1) mismatches (7) | 6 +1bp, -1bp, -860bp |
|  |  |  | 3 | 10329 | indel ≤ 5 bp (8) indel > 5 bp (1) mismatches (7) | 7 +1bp, -1bp, -802bp |
|  |  | Long-read | 1 | 11136 | indel ≤ 5 bp (17) mismatches (15) | 11 +1bp, 4 -1bp, 2 +2 bp (1 rep. G) |
|  |  |  | 2 | 11137 | indel ≤ 5 bp (17) mismatches (14) | 11 +1bp, +2bp, 4 -1bp, +3 bp (rep. G) |
|  |  |  | 3 | 11136 | indel ≤ 5 bp (17) mismatches (14) | 11 +1bp, 2 +2bp (1 rep. G), 4 -1bp |
|  |  | Hybrid | 1 | 11139 | indel ≤ 5 bp (16) mismatches (14) | 11 +1bp, +2bp, 3 -1bp, +4 bp (rep. G) |
|  |  |  | 2 | 11139 | indel ≤ 5 bp (17) mismatches (14) | 11 +1bp, +2bp, 4. -1bp, +5 bp (rep. G) |
|  |  |  | 3 | 11140 | indel ≤ 5 bp (16) mismatches (14) | 11 +1bp, +2bp, +5bp (rep. G), 3 -1bp |
| 3132 | 6331 | Short-read | 1 | 5467 | indel ≤ 5 bp (7) indel > 5 bp (1) mismatches (7) | 6 +1bp, -1bp, -869bp |
|  |  |  | 2 | 5550 | indel ≤ 5 bp (8) indel > 5 bp (1) mismatches (7) | 7 +1bp, -1bp, -787bp |
|  |  |  | 3 | 5383 | indel ≤ 5 bp (7) indel > 5 bp (1) mismatches (7) | 6 +1bp, -1bp, -953bp |
|  |  | Long-read | 1 | 6345 | indel ≤ 5 bp (16) mismatches (15) | 11 +1bp, +2bp, 3 -1bp, +4bp (rep. G) |
|  |  |  | 2 | 18620 | indel ≤ 5 bp (16) indel > 5 bp (1) mismatches (12) | 11 +1bp, 2 +2bp (1 rep. G), 3 -1bp, multiplicity: +12277 bp |
|  |  |  | 3 | 6346 | indel ≤ 5 bp (16) mismatches (14) | 11 +1bp, +3bp, 3 -1bp, +4 bp (rep. G) |
|  |  | Hybrid | 1 | 6342 | indel ≤ 5 bp (14) mismatches (14) | 11 +1bp, 2 +2bp (1 rep. C), 4 -1bp |
|  |  |  | 2 | 6350 | indel ≤ 5 bp (15) indel > 5 bp (1) mismatches (15) | +9bp (rep. C), 11 +1bp, +2bp, 3 -1bp |
| 3127 | 7452 | Short-read | 1 | 6652 | indel ≤ 5 bp (1) indel > 5 bp (1) | +1bp (rep. C), -801bp |
|  |  |  | 2 | 6586 | indel > 5 bp (1) | -866bp |
|  |  | Long-read | 1 | 7464 | indel ≤ 5 bp (5) indel > 5 bp (1) mismatches (7) | +7bp, +3bp, 3 +1bp, -1bp |
|  |  |  | 2 | 7457 | indel ≤ 5 bp (12) mismatches (7) | +4bp (rep. G), +2bp, 5 +1bp, 4 -1bp, -2bp |
|  |  |  | 3 | 7463 | indel ≤ 5 bp (12) mismatches (7) | +5bp (rep. G), +2bp, 7 +1bp, 3 -1bp |
|  |  | Hybrid | 1 | 7462 | indel ≤ 5 bp (9) indel > 5 bp (1) mismatches (7) | +6bp (rep. C), 5 +1bp, +2bp, 3 -1bp |
|  |  |  | 2 | 7459 | indel ≤ 5 bp (9) mismatches (7) | 5 +1bp, 2 +2bp (1 rep. G), 2 -1bp |

**Table S6. Summary of results from reference-based assemblies.**

| **Plasmid ID** | **Expected Reference Size (bp)** | ***De Novo* Assembly Size (bp)** | **Reference-Based Assembly Size [with Sanger] (bp)** |
| --- | --- | --- | --- |
| 3121 | 7784 | 7765 | 8484 |
| 3123 | 6828 | 6832 | 7535 |
| 3124 | 6817 | 6819 | 7674 |
| 3134 | 15045 | 15044 | 15892 |
| 3127 | 7452 | 7459* | 8010 [8317] |
| 3130 | 11125 | 11139* | 12578 [11908] |
| 3131 | 6064 | 6078* | 6749 [6722] |
| 3132 | 6331 | 6342* | 7104 [7237] |
| 3133 | 14194 | 14191 | 14785 [14785] |

* = Hybrid assembly size

**Table S7. Cost analysis of 96 samples on each sequencing technology (prices in USD as of December 2023).**

| **Sequencing Method** | **Consumables Costs** | **Instrument Costs** | **Labor Costs** | **Total Cost** | **Cost per sample** |
| --- | --- | --- | --- | --- | --- |
| **MinION** | $551.60 | $6.15 | $139.49 | $697.21 | $7.26 |
| **Sanger** | $1,259.31 | $286.01 | $69.74 | $1,615.06 | $16.82 |
| **MiSeq** | $4,354.49 | $364.11 | 185.99 | $4,904.59 | $51.09 |

_____________________________________________________________________________________________

**Supplementary Figures**

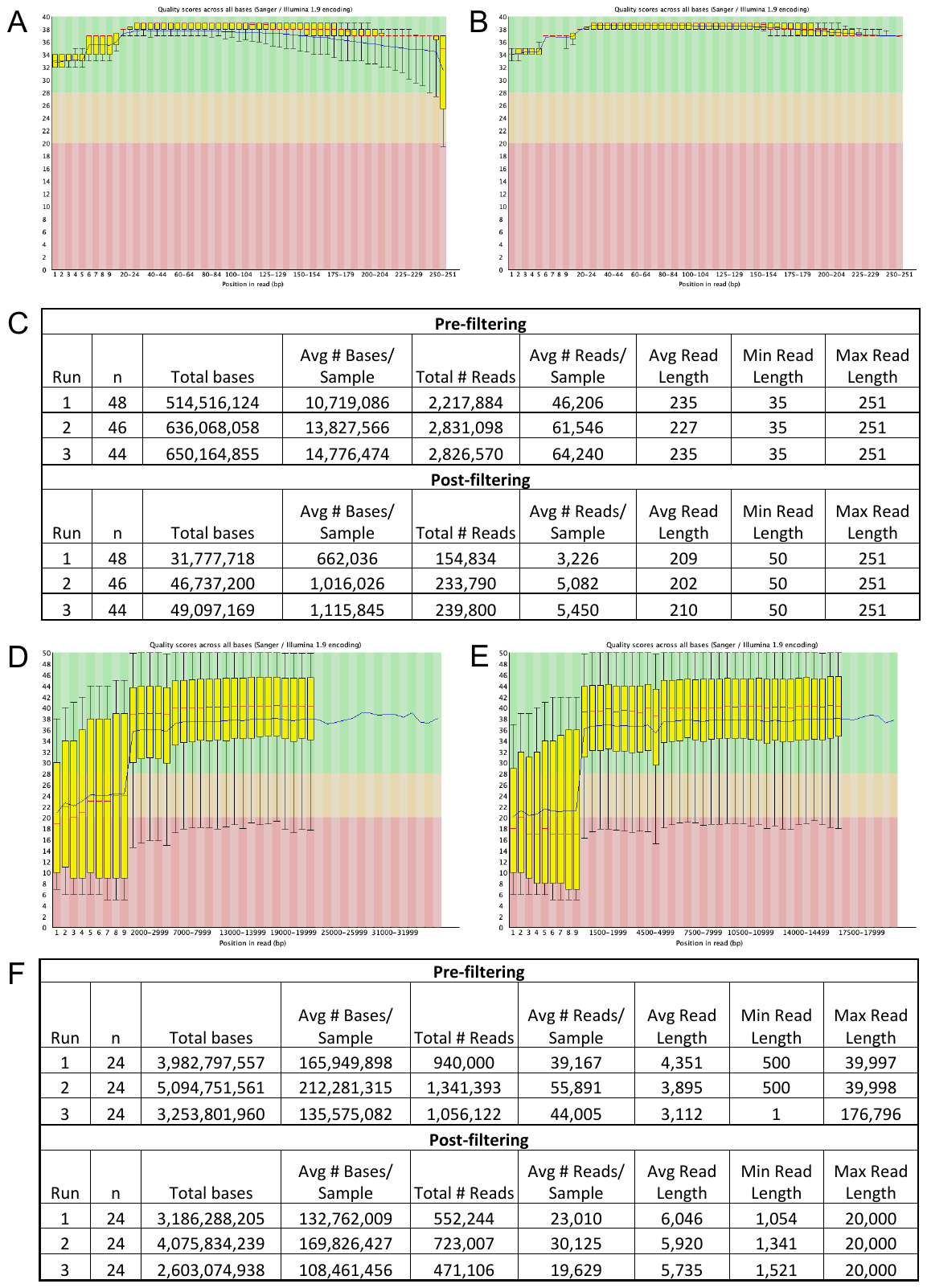

**Figure S1. Summary of sequencing read statistics before and after filtering.** Representative short read data quality (A) before and (B) after filtering. (C) Short reads were trimmed and filtered using Trimmomatic. Representative long read data quality (D) before and (E) after filtering. (F) Long reads were trimmed and filtered using Filtlong.

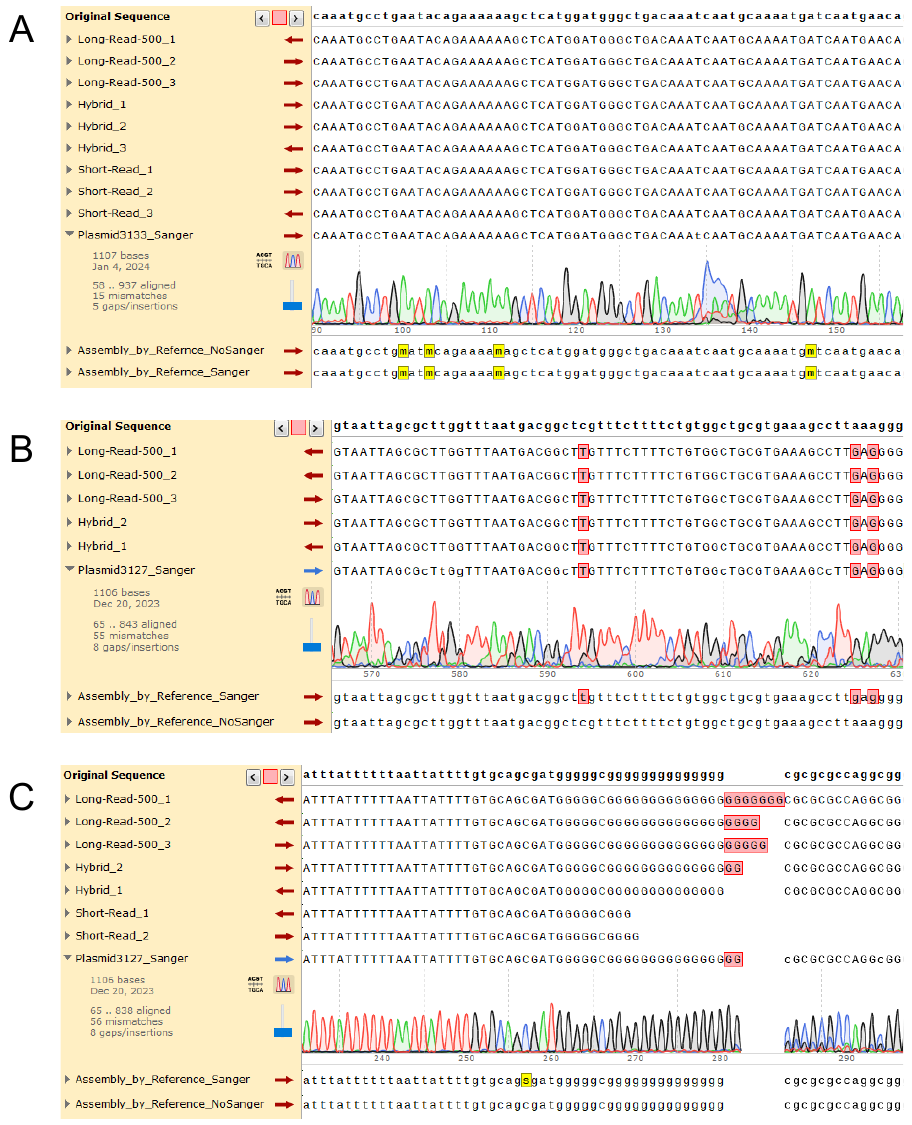

**Figure S2. Reference-based assemblies are less accurate than *de novo* assemblies.**

The sequences of reference-based assemblies and *de novo* short-read, long-read, and hybrid assemblies were aligned to the reference sequence (top, “Original Sequence”). (A) Representative reference-based assembly containing non-standard nucleotides despite clean Sanger data. (B) Representative example where Sanger data improves reference-based assembly and supports the *de novo* assembly sequences. (C) Representative example of reference-based assembly demonstrating reference bias, even when Sanger data is provided.

_____________________________________________________________________________________________

**Supplementary References**

1.	Chevreux, B., Wetter, T. and Suhai, S. (1999) Genome Sequence Assembly Using Trace Signals and Additional Sequence Information. In *German Conference on Bioinformatics.* ***99,*** *1, 45-56.*

2.	Cock, P.J., Grüning, B.A., Paszkiewicz, K. and Pritchard, L. (2013) Galaxy tools and workflows for sequence analysis with applications in molecular plant pathology. *PeerJ*, **1**, e167.

3.	The Galaxy Community. (2022) The Galaxy platform for accessible, reproducible and collaborative biomedical analyses: 2022 update. *Nucleic Acids Research*, **50**, W345-W351.

4.	Wilson, M.L., Cai, Y., Hanlon, R., Taylor, S., Chevreux, B., Setubal, J.C., Tyler, B.M. and Peccoud, J. (2012) Sequence verification of synthetic DNA by assembly of sequencing reads. *Nucleic Acids Research*, **41**, e25-e25.
